# RBM6-mediated alternative splicing facilitates the adaptation of Eurasian avian-like H1N1 swine influenza virus

**DOI:** 10.64898/2026.07.22.740191

**Authors:** Jiahui Zou, Shaoyu Tu, Huimin Sun, Chuhan Xiong, Meijun Jiang, Jinli Guo, Shuang Tang, Tong Chen, Thomas P Peacock, Wen Su, Wendy S Barclay, Hongbo Zhou

## Abstract

The Eurasian avian□like (EA) H1N1 swine influenza virus (SIV), derived from avian influenza viruses (AIV), poses a serious threat to public health due to its capacity for cross□species transmission and pandemic emergence. The molecular determinants underlying its replication advantage over AIV remain poorly defined. Here, we identify RNA□binding motif protein 6 (RBM6) as a novel host factor that differentially regulates the replication of EA H1N1 SIV and AIV. Mechanistically, RBM6 binds to the critical M901 site of the viral M segment, thereby modulating RNA splicing. Substitution of M901C with M901T markedly reduced RBM6 binding, impaired M segment splicing, and attenuated viral replication both *in vitro* and *in vivo*. Conservation analysis revealed that M901T is common in avian strains, whereas M901C is predominantly maintained in swine strains, underscoring M901C as a determinant of swine adaptation. Complementation experiments further demonstrated that swine RBM6, but not avian RBM6, restored EA H1N1 SIV replication. Taken together, our findings uncover a previously unrecognized role of RBM6 in shaping influenza virus replication and highlight the RBM6-M901C axis may serve as potential targets for controlling influenza virus adaptation and interspecies transmission.

## Introduction

Influenza A virus (IAV) is a major global health threat, responsible for recurrent epidemics and occasional pandemics ^1,2^. Avian influenza viruses (AIVs) are of particular concern because their ability to cross species barriers and infect mammals can result in severe disease and even human fatalities ^3,4^. Pigs, which express both avian□type α□2,3 and human□type α□2,6 sialic acid (SA) receptors, function as important intermediate hosts that facilitate the adaptation of AIVs to mammalian systems ^5^. The Eurasian avian□like (EA) H1N1 swine influenza viruses (SIVs), derived from the avian strain A/duck/Bavaria/2/1979, have become established in pig populations across Europe and Asia and have occasionally infected humans ^6,7^. During their evolution, EA H1N1 SIVs shifted from exclusive binding to α□2,3 SA to dual recognition of α□2,3 and α□2,6 SA, with recent isolates showing increased affinity for α□2,6 SA, a hallmark of mammalian adaptation ^7,8^. Following the 2009 H1N1 pandemic, reverse zoonosis reintroduced the virus into pigs, producing reassortants between EA H1N1 SIV and H1N1pdm09 ^9,10^. Some of these reassortants demonstrated efficient respiratory droplet transmission in ferrets, underscoring their pandemic potential ^11^. EA H1N1 SIV, therefore, represents a valuable model for investigating the mechanisms of avian influenza interspecies transmission and assessing its implications for public health.

Host factors are fundamental to the regulation of IAV interspecies transmission, as they act at multiple stages of the viral life cycle, including entry, replication, splicing, assembly, and release ^12–17^. These interactions determine the efficiency of infection in different hosts. Among the diverse mechanisms involved, the regulation of viral RNA splicing has emerged as a particularly critical determinant of host adaptation ^13,14^. Splicing ensures the balanced production of essential viral proteins, and even subtle differences in host splicing machinery can either restrict or facilitate viral replication across species barriers ^18–20^. Heterogeneous nuclear ribonucleoprotein M (hnRNPM) displays species-specific activity by modulating viral mRNA splicing, thereby influencing replication efficiency in avian and mammalian cells ^13^. Transformer-2 protein homolog alpha (TRA2A) binds splicing silencer motifs within viral transcripts, limiting replication of avian influenza viruses in mammals while supporting replication of human-adapted strains ^14^. These findings emphasize RNA splicing as a central mechanism in shaping viral host range. Furthermore, they suggest that additional RNA-binding proteins with splicing regulatory functions remain to be identified, and their characterization will be essential for a comprehensive understanding of the molecular basis of influenza interspecies transmission.

In this study, we identify RNA□binding motif protein 6 (RBM6) as a novel host factor that regulates influenza virus replication in a subtype□specific manner. RBM6 interacts with the M901 site of the viral M segment to influence RNA splicing, and the presence of M901C enhances replication efficiency in swine. Conservation analysis shows that M901T predominates in avian strains, while M901C is characteristic of swine strains, marking M901C as a determinant of swine adaptation. Complementation assays further demonstrate that swine RBM6, but not avian RBM6, restore EA H1N1 SIV replication. Collectively, these findings establish swine RBM6 as a critical regulator of influenza virus replication dynamics and highlight the RBM6-M901C axis as a potential target for controlling viral adaptation and interspecies transmission.

## Results

### RBM6 is identified as an EA H1N1 SIV host-dependent factor in porcine cells

RBM6 was identified as an necessary host factor by genome-wide CRISPR screening ^15,21^. To investigate the role of RBM6 in IAV infection, we generated RBM6□knockout (RBM6-KO) porcine kidney□15 (PK□15) cells using CRISPR/Cas9 technology, as confirmed by the absence of RBM6 protein expression and genomic base deletion (Supplementary Fig. 1a-b). Notably, RBM6 deficiency did not affect cell viability (Supplementary Fig. 1c). It was revealed that knockout of RBM6 inhibited the replication of avian-origin A/duck/Bavaria/2/1997 (Bav/H1N1) (Fig. 1a). We further evaluated the role of RBM6 in the infection of different EA H1N1 SIV strains and observed significantly reduced virus titers in RBM6-deficient cells infected with swine-origin A/Swine/HuBei/221/2016 (HuB/H1N1) and human-isolated A/Hunan/42443/2015 (HuN/H1N1) EA H1N1 SIV (Fig. 1b-c). Furthermore, the inhibitory effect of RBM6-deficient cells was more pronounced for EA H1N1 SIV (HuB/H1N1 and HuN/H1N1) than for avian-origin (Bav/H1N1) strain (Fig. 1a-c). Further investigation into the role of avian RBM6 in AIV replication revealed that it exerted no significant effect on the replication of AIV (Bav/H1N1) (Fig. 1d), suggesting that species□specific RBM6 is involved in the interspecies transmission of influenza A viruses.

**Figure 1.**
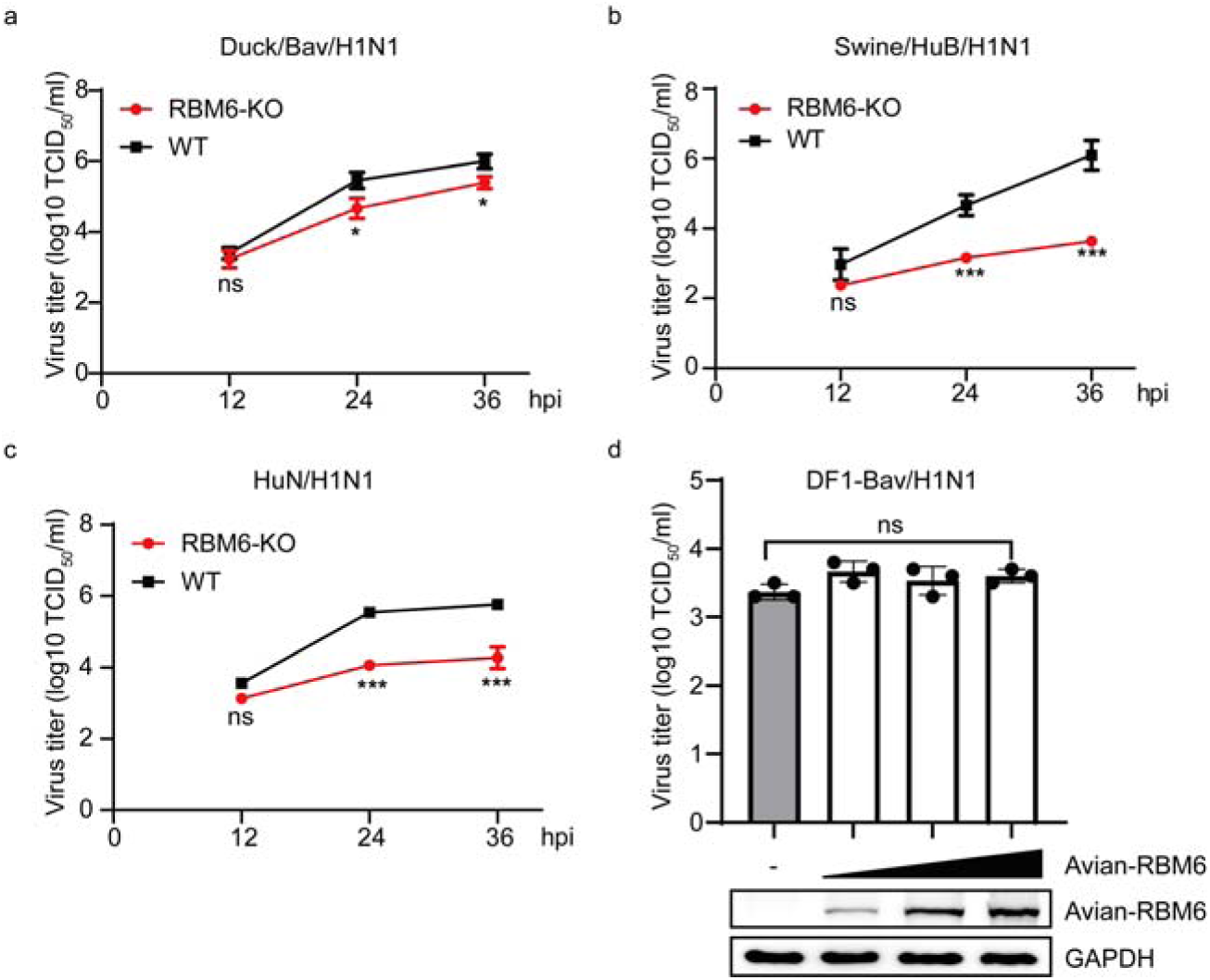
RBM6 is identified as an EA H1N1 SIV host-dependent factor in porcine cells. (a-c) Effects of RBM6 knockout in PK-15 on the replication of (a) Bav/H1N1, (b) HuB/H1N1, and (c) HuN/H1N1. RBM6-KO and WT PK-15 cells were infected IAV strains at a MOI of 0.01, and the viral titers at indicated timepoints were determined by TCID_50_. (d) Effect of avian RBM6 overexpression on AIV replication. DF-1 cells were transfected with increasing amounts of exogenous avian HA-RBM6. At 24 h post-transfection, the cells were infected with Bav/H1N1 at a MOI of 0.1. Viral titers were measured at 24 h post-infection using the TCID_50_ assay. Error bar in panels a-d indicates the standard deviation. The data shown in panels a-d are means□±□SD (n□=□3 for biologically independent experiments). Statistical analysis was performed using unpaired, two-tailed Student’s *t*-test. (ns, *P* > 0.05; *, *P* < 0.05; ***, *P* < 0.001).

### RBM6 facilitates the viral splicing

To determine the stage of the EA H1N1 SIV infection cycle at which RBM6 is involved, we examined the cellular distribution of viral nucleoprotein (NP) in infected WT and RBM6□KO cells. Up to 6 hpi, no differences in NP expression or localization were detected, and NP had translocated into the nucleus in both cell types. By 9 hpi, however, RBM6 deficiency led to reduced NP expression (Supplementary Fig. 2), suggesting that RBM6 supports viral replication after nuclear import.

Because the homologous genes of RBM6, RBM5 and RBM10, have been reported to participate in RNA splicing ^22–24^, we hypothesized that RBM6 may also be involved in the splicing of both host and viral transcripts, thereby regulating viral replication. To elucidate the role of RBM6 in IAV RNA splicing, RBM6-KO and WT cells were infected with the Bav/H1N1 strain, and the splicing of viral M and NS transcripts were analyzed at both RNA and protein levels. RBM6 deficiency had no discernible effect on the splicing of M and NS segments in Bav/H1N1□infected cells (Fig. 2a-c). We next assessed the function of RBM6 during RNA splicing of EA H1N1 SIV, RBM6-KO and WT cells were infected with the swine-origin HuB/H1N1 and human-isolated HuN/H1N1 EA H1N1 SIV. In contrast to avian□origin Bav/H1N1, RBM6 depletion significantly impaired the splicing of M and NS segments in both EA H1N1 SIV strains, with this inhibitory effect being notably stronger than that observed in the avian□origin virus (Fig. 2d–i). Notably, this gradient of splicing inhibition paralleled the impact of RBM6 knockout on viral replication, thereby supporting that RBM6 regulates IAV replication through modulation of viral RNA splicing.

**Figure 2.**
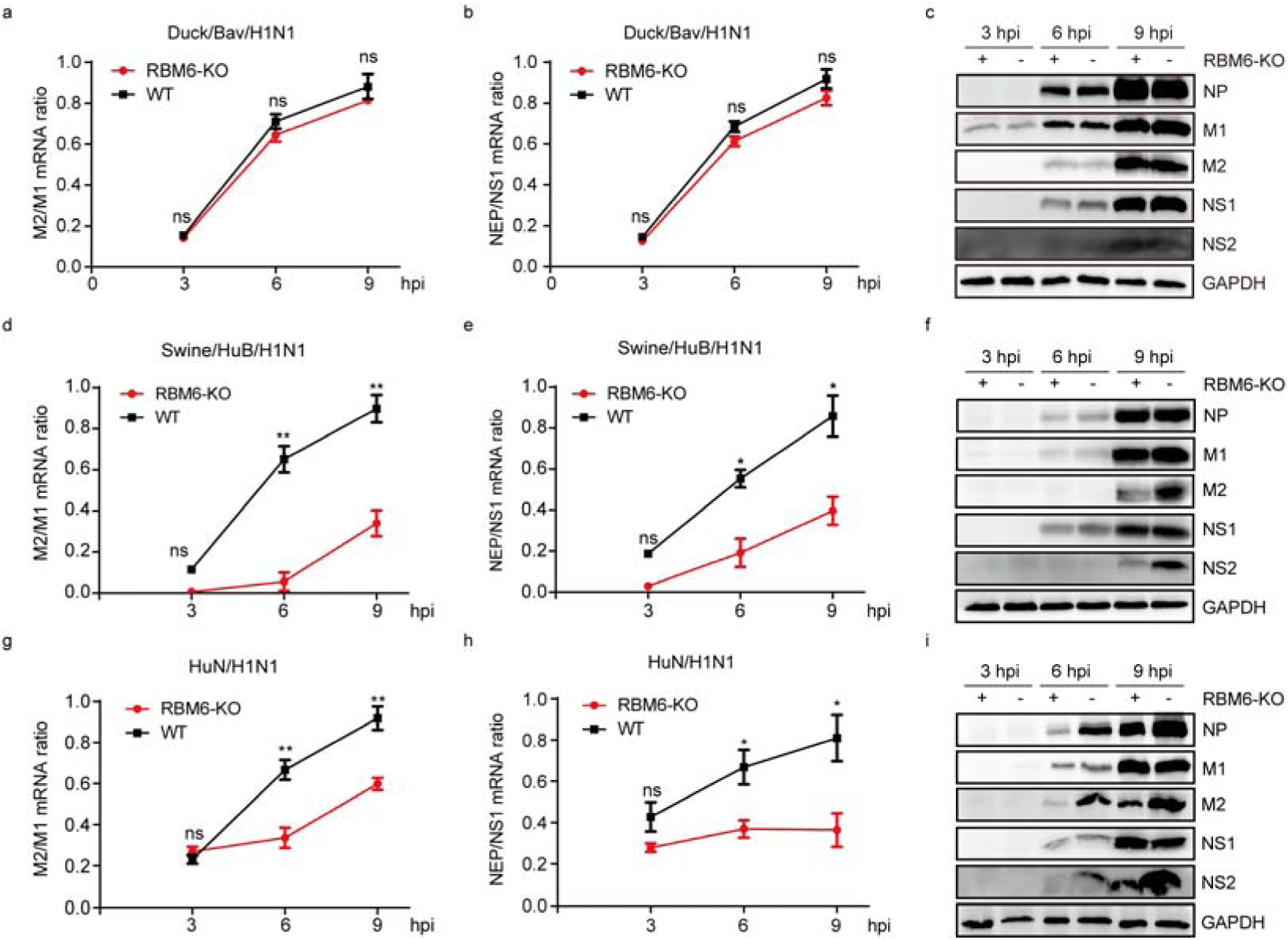
RBM6 facilitated the splicing of EA H1N1 SIV. Wild-type and RBM6-KO PK-15 cells were respectively infected with (a-c) Bav/H1N1, (d-f) HuB/H1N1, and (g-i) HuN/H1N1 for 3, 6, and 9 h. Cells were harvested for total RNA isolation or western blot analysis. Total RNA was extracted, and the mRNAs of viral M1, M2, NS1, and NS2 were quantified by qRT-PCR. Cell lysates were analyzed by western blot to assess the expression of viral M1, M2, NS1, NS2, and NP, with GAPDH used as a loading control. Error bar in panels a-b, d-e, and g-h indicates the standard deviation. The data shown in panels a-b, d-e, and g-h are means□±□SD (n□=□3 biologically independent experiments). Statistical analysis was performed using unpaired, two-tailed Student’s *t*-test. (ns, *P* > 0.05; *, *P* < 0.05; **, *P* < 0.01).

### RBM6 regulates IAV RNA splicing and replication through the critical M901 site on the M segment

Given that RBM6 regulates IAV replication through modulation of M and NS splicing, we next sought to identify the binding sites of RBM6 on viral M and NS transcripts. A stable cell line expressing halo-tagged RBM6 was established, and CLIP-Seq analysis was performed to characterize RBM6-binding motifs (Fig. 3a) ^25^. This approach yielded multiple candidate motifs (Supplementary Fig. 3), among which the top-ranked and overlapped motifs were selected for further validation. RNA pull-down assays confirmed that M-7-3 and NS-8-5 exhibited the strongest binding affinity to RBM6 (Fig. 3b-c). Consistently, the M-7-3 motif showed substantially higher binding capacity than NS-8-5 (Fig. 3d). Based on these findings, subsequent investigations focused primarily on the M segment.

**Figure 3.**
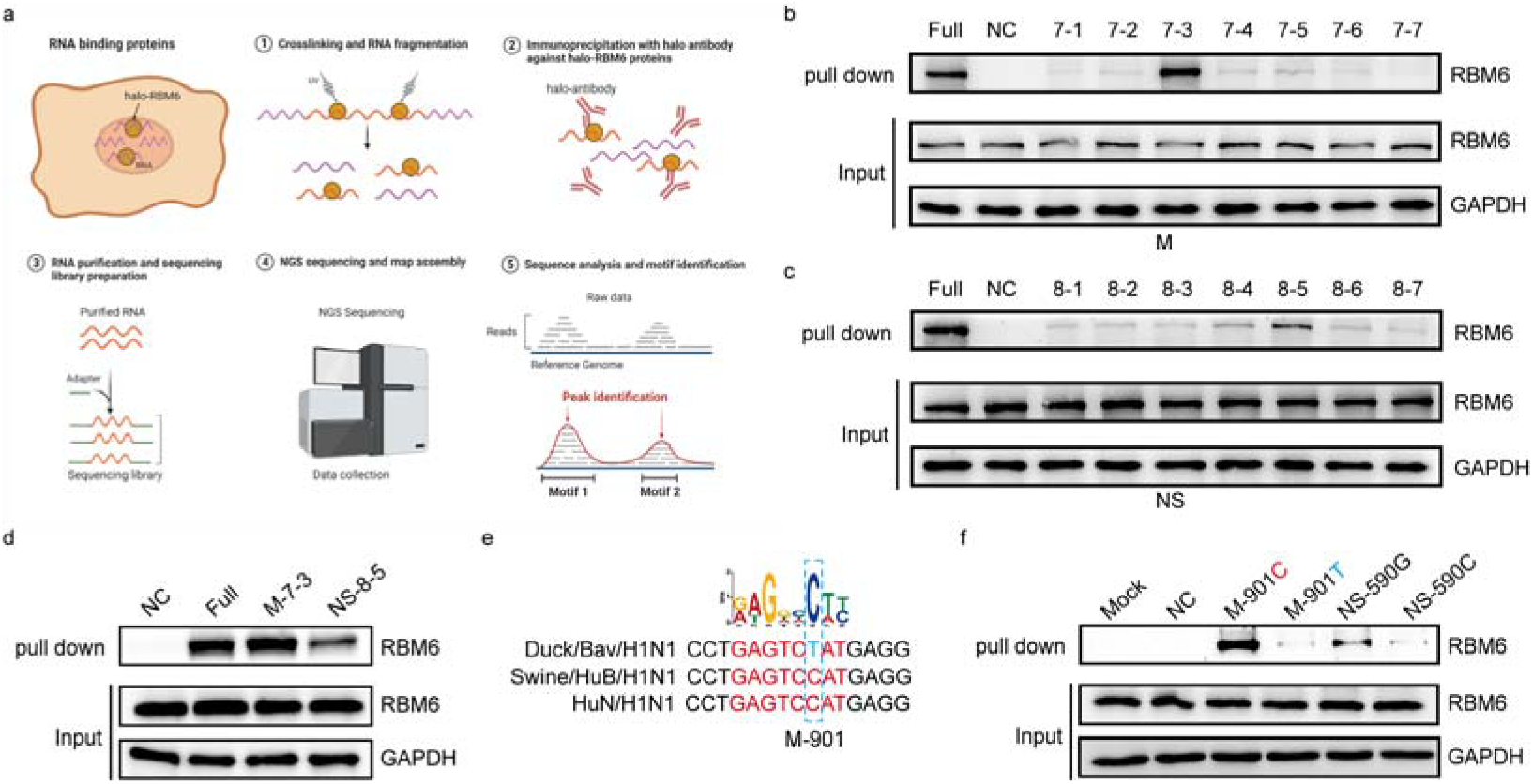
Viral M901C is critical for binding to RBM6. (a) Schematic model for screening RBM6□binding motifs on viral M and NS segments. PK□15 cells stably expressing Halo□tagged RBM6 were infected with HuB/H1N1 at an MOI of 5 for 9 h, followed by UV irradiation (365 nm) to induce covalent crosslinking between RBM6 and its bound RNAs. Cell lysates were then analyzed by CLIP□seq. (b–c) Validation of RBM6□binding motifs identified on the viral (b) M and (c) NS segments. Selected motifs were chemically synthesized, biotin□labeled, and their binding activities to RBM6 were evaluated using RNA pull down assays. (d) Comparison of the binding activities of M and NS motifs to RBM6. Chemically synthesized and biotin□labeled motifs were tested by RNA pull down assay. (e) Sequence comparison of M motifs from different viruses. (f) Effect of mutations introduced into full□length M or NS on RBM6 binding. Full□length M and NS constructs carrying the M C901T or NS G590C mutations were generated by site□directed mutagenesis and *in vitro* transcription, and their binding abilities to RBM6 were evaluated using RNA pull down assays.

Because RBM6 knockout exerted differential effects on splicing and replication of IAVs from distinct species, we hypothesized that sequence variation in viral M motifs may underlie these differences. Comparative sequence analysis revealed difference at position M901 among IAVs of different origins (Fig. 3e). To test the functional relevance of this site, mutations were introduced into the M901 within the full-length M segment. The results showed that mutation of M901 markedly reduced the binding capacity to RBM6 (Fig. 3f), indicating that M901 represents a critical determinant for RBM6 interaction. These findings suggest that variation at M901 contributes to the species-specific effects of RBM6 on IAV RNA splicing and replication.

### Viral M901C determined the replication of EA H1N1 SIV

To investigate the impact of the M C901T mutation on viral replication and splicing in EA H1N1 SIV, we introduced this substitution into the HuN/H1N1 strain to generate the mutant virus. We first compared the ability of wild□type and M C901T mutant HuN/H1N1 to associate with RBM6, and found that the mutation markedly reduced RBM6 binding (Fig. 4a). In addition, the M C901T mutant exhibited decreased splicing of the M segment, while NS splicing remained unaffected (Fig. 4b–d). Replication efficiency was then assessed by measuring viral titers over time. The M C901T mutant displayed attenuated replication compared with wild□type HuN/H1N1 (Fig. 4e). To further clarify the role of RBM6 in regulating M901, wild□type and mutant viruses were tested in both WT and RBM6 KO PK 15 cells. Growth curve analysis revealed that wild□type virus replicated to significantly higher titers in WT cells, whereas replication in RBM6□KO cells was comparable to that of the mutant virus (Fig. 4f). Together, these results demonstrate that the M C901T mutation impairs both viral replication and M segment splicing in EA H1N1 SIV, highlighting the critical role of the M901 site in RBM6□mediated regulation of influenza virus replication.

**Figure 4.**
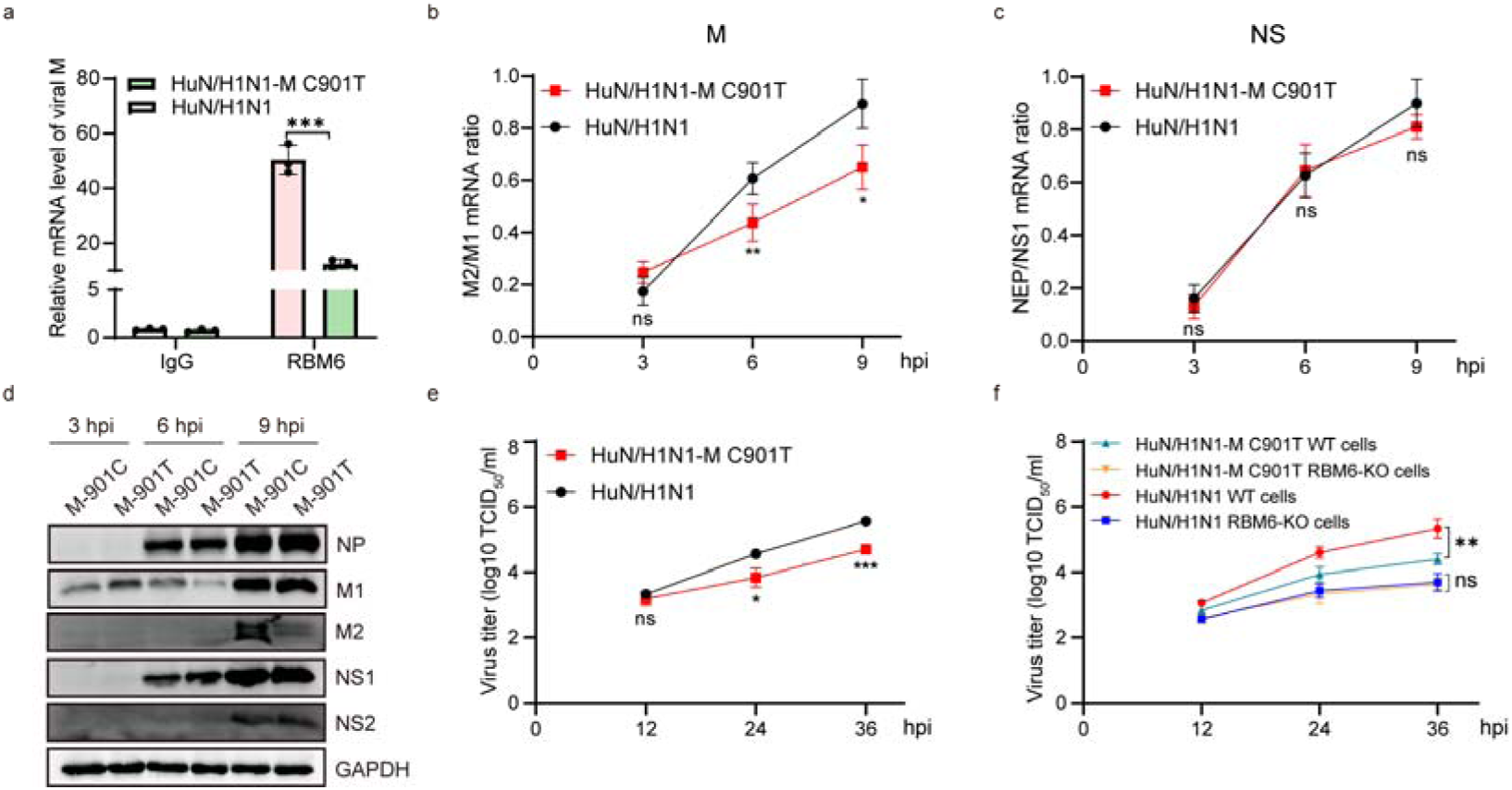
Viral M901C determined the replication of EA H1N1 SIV. (a) Binding activities of wild□type or M C901T mutant HuN/H1N1 viral M to RBM6. PK□15 cells were infected with wild type or M C901T mutant HuN/H1N1 for 12 h. Anti□RBM6 or control IgG antibodies were used to immunoprecipitate bound RNA, followed by total RNA extraction and qRT□PCR analysis of viral M. (b–d) Effects of the M C901T mutation on splicing in EA H1N1 SIV. PK□15 cells were infected with wild□type or M C901T mutant HuN/H1N1 for 3, 6, and 9 h. Cells were harvested for total RNA isolation or western blot analysis. (b–c) Viral M1, M2, NS1, and NS2 mRNAs were quantified by qRT□PCR. (d) Protein expression of M1, M2, NS1, NS2, and NP was assessed by western blot, with GAPDH as a loading control. (e) Effects of the M C901T mutation on EA H1N1 SIV replication. PK 15 cells were infected with wild□type or M C901T mutant HuN/H1N1 at an MOI of 0.01, and viral titers at the indicated time points were determined by TCID_50_. (f) Growth kinetics of wild□type and M C901T mutant HuN/H1N1 in WT and RBM6□KO PK□15 cells. Cells were infected at MOI = 0.01, and viral titers at the indicated time points were determined by TCID_50_. Error bars in panels a-c and e-f indicates the standard deviation. The data shown in panel a-c and e-f is means□±□SD (n□=□3 biologically independent experiments). Statistical analysis was performed using unpaired, two-tailed Student’s *t*-test. (ns, *P* > 0.05; *, *P* < 0.05; **, *P* < 0.01; ***, *P* < 0.001).

### Swine RBM6 promotes the adaptation of EA H1N1 SIV M901C

EA H1N1 SIVs originated from Eurasian avian H1N1 ^6,26^. Although host factors and viral determinants contributing to adaptive evolution have been identified ^9,27,28^, the role of the M segment in this process remains poorly defined. Conservation analysis of the M901 site in avian H1N1 and EA H1N1 SIV revealed that M901C was predominantly conserved in EA H1N1 SIV, whereas M901T was more frequently observed in avian H1N1 (Fig. 5a). Notably, in the early stage of EA H1N1 SIV, the M901T variant was present but subsequently underwent adaptive mutation to M901C (Fig. 5b), suggesting that M901C is a key determinant of EA H1N1 SIV evolutionary adaptation. Further analysis of M901 conservation across influenza viruses demonstrated that M901T was more common in AIVs, while M901C was predominantly maintained in SIVs (Fig. 5c-d). Collectively, these findings indicate that M901C plays a critical role in the adaptation of AIV to swine. Moreover, in RBM6-deficient PK-15 cells, complementation with swine RBM6 successfully restored viral replication, whereas avian RBM6 was unable to restore replication (Fig. 5e). These results demonstrate that RBM6 from swine, but not avian RBM6, promotes the adaptation of EA H1N1 SIV M901C.

**Figure 5.**
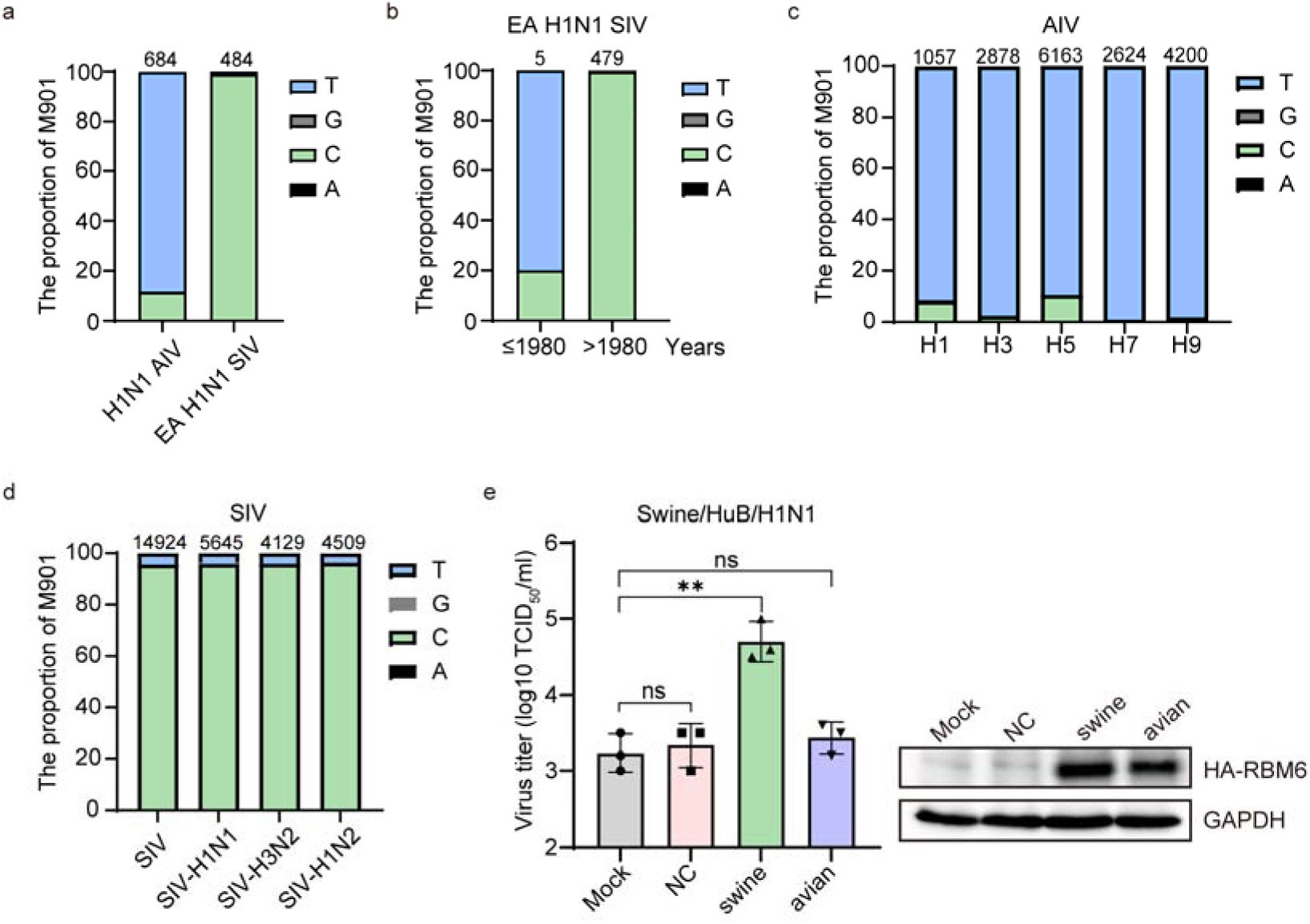
Swine RBM6 promotes the adaptation of EA H1N1 SIV M901C. (a) Conservation analysis of the M901 site in H1N1 AIV and EA H1N1 SIV. (b) Temporal conservation analysis of the M901 site in EA H1N1 SIV. (c–d) Conservation analysis of the M901 site in (c) AIVs and (d) SIVs strains downloaded from the National Center for Biotechnology Information (NCBI). (e) Restoration of swine RBM6 promotes replication of EA H1N1 SIV. Exogenous RBM6 (HA□RBM6) from swine and avian species was transfected into RBM6□KO PK□15 cells. Transfected cells were infected with HuB/H1N1 at an MOI of 0.1, and viral titers at 24 hpi were determined by TCID_50_. Corresponding protein expression was detected by western blot. Error bar indicates the standard deviation. The data shown in panel e are means□±□SD (n□=□3 biologically independent experiments). Statistical analysis was performed using unpaired, two-tailed Student’s *t*-test. (ns, *P* > 0.05; **, *P* < 0.01; ***, *P* < 0.001).

### RBM6 and viral M 901C promoted the infection of EA H1N1 SIV *in vivo*

To further investigate the role of RBM6 and the M C901T mutation in EA H1N1 SIV replication *in vivo*, wild-type and RBM6-deficient mice were respectively challenged with wild-type or M C901T mutant HuN/H1N1 virus (Fig. 6a). Mouse weight loss and survival were monitored daily for 14 days post-challenge. RBM6-deficient mice, as well as mice infected with the mutant M C901T virus, exhibited attenuated infection, characterized by reduced weight loss and enhanced survival relative to wild-type mice or those challenged with the HuN/H1N1 virus (Fig. 6b-c). Notably, RBM6-deficient mice infected with the mutant M C901T virus exhibited modestly reduced weight loss and achieved a 90% survival rate (Fig. 6b-c). We also observed a significant decrease in viral titers in the lungs of RBM6-deficient mice and in those challenged with the mutant M C901T virus (Fig. 6d-e). Histopathological analysis of the lungs of wild-type mice infected with the HuN/H1N1 virus revealed moderate to severe bronchiolar necrosis, pulmonary oedema, and extensive inflammatory cell infiltration (Fig. 6f). In contrast, examination of the lungs from RBM6-deficient mice and those challenged with the M C901T mutant HuN/H1N1 virus demonstrated a marked reduction in lymphoid tissue infiltration relative to control animals (Fig. 6f). Meanwhile, weaker viral nucleoprotein (NP) antigen signals were detected in the lungs of RBM6-deficient mice and in those challenged with the M C901T mutant virus (Fig. 6g). These results demonstrated that RBM6 deficiency and the M C901T mutation confer significant protection against EA H1N1 SIV challenge *in vivo*.

**Figure 6.**
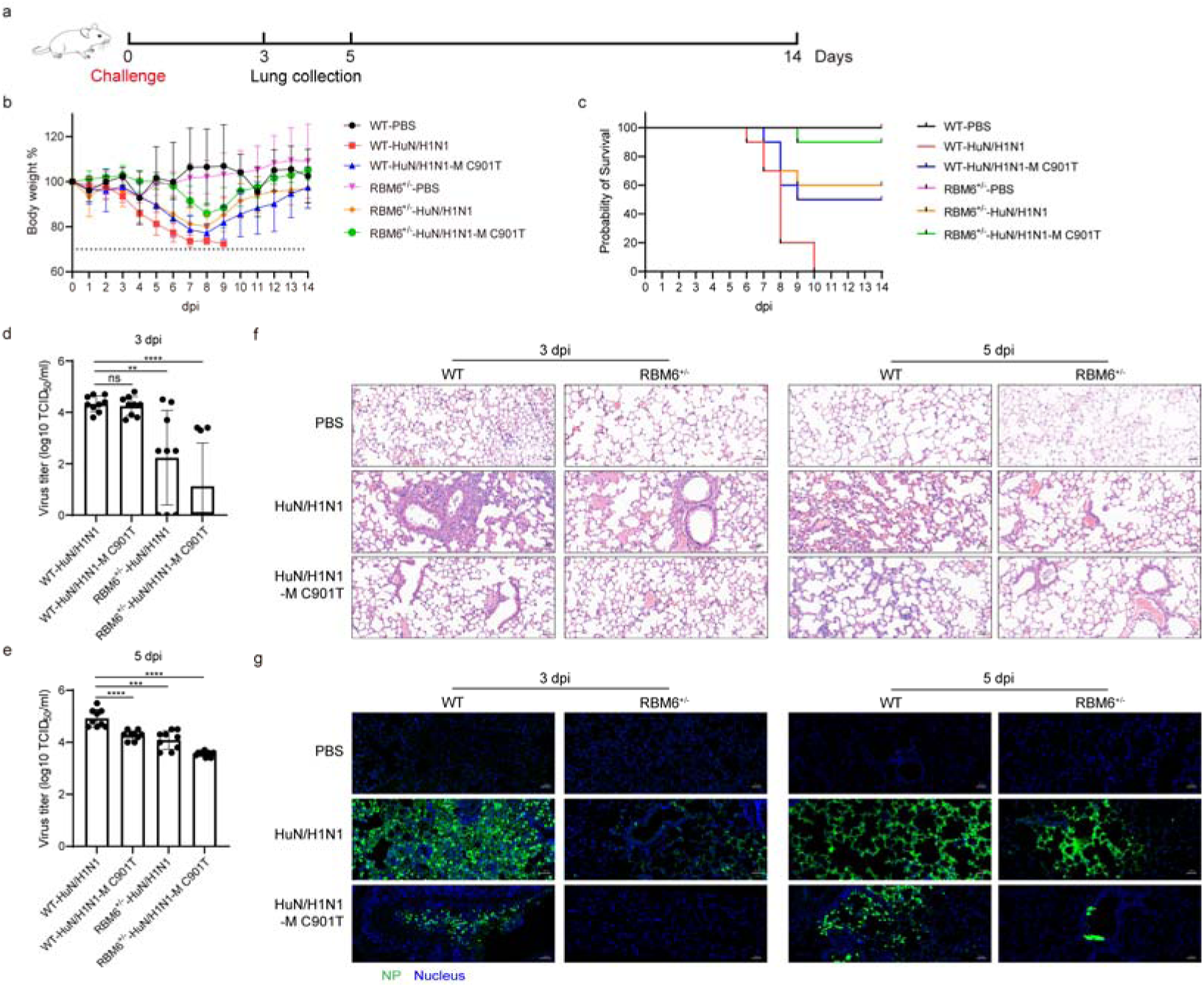
RBM6 and viral M901C promoted the infection of EA H1N1 SIV *in vivo*. (a) Schematic of HuN/H1N1 strain challenge in an experimental mouse model. Wild-type or RBM6 deficiency mice were challenged with wild-type or M C901T mutant HuN/H1N1, and monitored for 14 days. (b-c) Weight loss and Mortality of wild-type and M C901T mutant HuN/H1N1 strain infected wild-type and RBM6 deficiency mice. Mice with a body weight loss of more than 30% were euthanized according to the ethical principles of animal welfare. Each treatment group had ten mice (n□=□10) per group. (d-e) Virus titers in the lungs of infected mice (d) 3 days and (e) 5 days after infection. Each treatment group had three mice (n□=□3) per group. (f) Hematoxylin and eosin (H&E) staining of pathological lesions in the lungs of wild-type and M C901T mutant HuN/H1N1 strain infected wild-type and RBM6 deficiency mice at 3 and 5 days post-challenge. Scale bars, 50 μm. (g) Immunofluorescence staining of lung sections from wild-type and M C901T mutant HuN/H1N1 strain infected wild-type and RBM6 deficiency mice at 3 and 5 days post-challenge. The viral NP antigen was stained green, and the nucleus was stained blue. Scale bars, 50 μm. The images in panels f and g are representative of three independent experiments. Error bar in panels b, d, and e indicates the standard deviation. The data shown in panels b, d, and e are means□±□SD (n = 10 for b and n□=□3 for d and e biologically independent experiments). Statistical analysis was performed using unpaired, two-tailed Student’s *t*-test. (ns, *P* > 0.05; **, *P* < 0.01; ***, *P* < 0.001; ****, *P* < 0.0001).

## Discussion

The EA H1N1 SIVs originated from H1N1 AIVs, yet the molecular determinants underlying their interspecies transmission remain incompletely defined. In this study, we identified RBM6 as a novel host factor that differentially regulates the replication of EA H1N1 SIV and AIV. Mechanistically, swine RBM6 interacts with the critical M901 site of the viral M segment, thereby modulating RNA splicing and enhancing viral gene expression. Conservation analysis revealed that the M901T is common in avian strains, whereas M901C variant is predominantly maintained in swine strains, underscoring M901C as a molecular signature of swine adaptation. Importantly, the acquisition of M901C increased compatibility with swine RBM6, resulting in stronger RBM6 binding, enhanced M segment splicing, and improved viral replication efficiency in swine cells (Fig. 7). Collectively, RBM6 plays a pivotal role in coordinating host-virus interactions at the M901 site, thereby fostering efficient replication in swine and promoting the evolutionary adaptation of EA H1N1 SIV.

**Figure 7.**
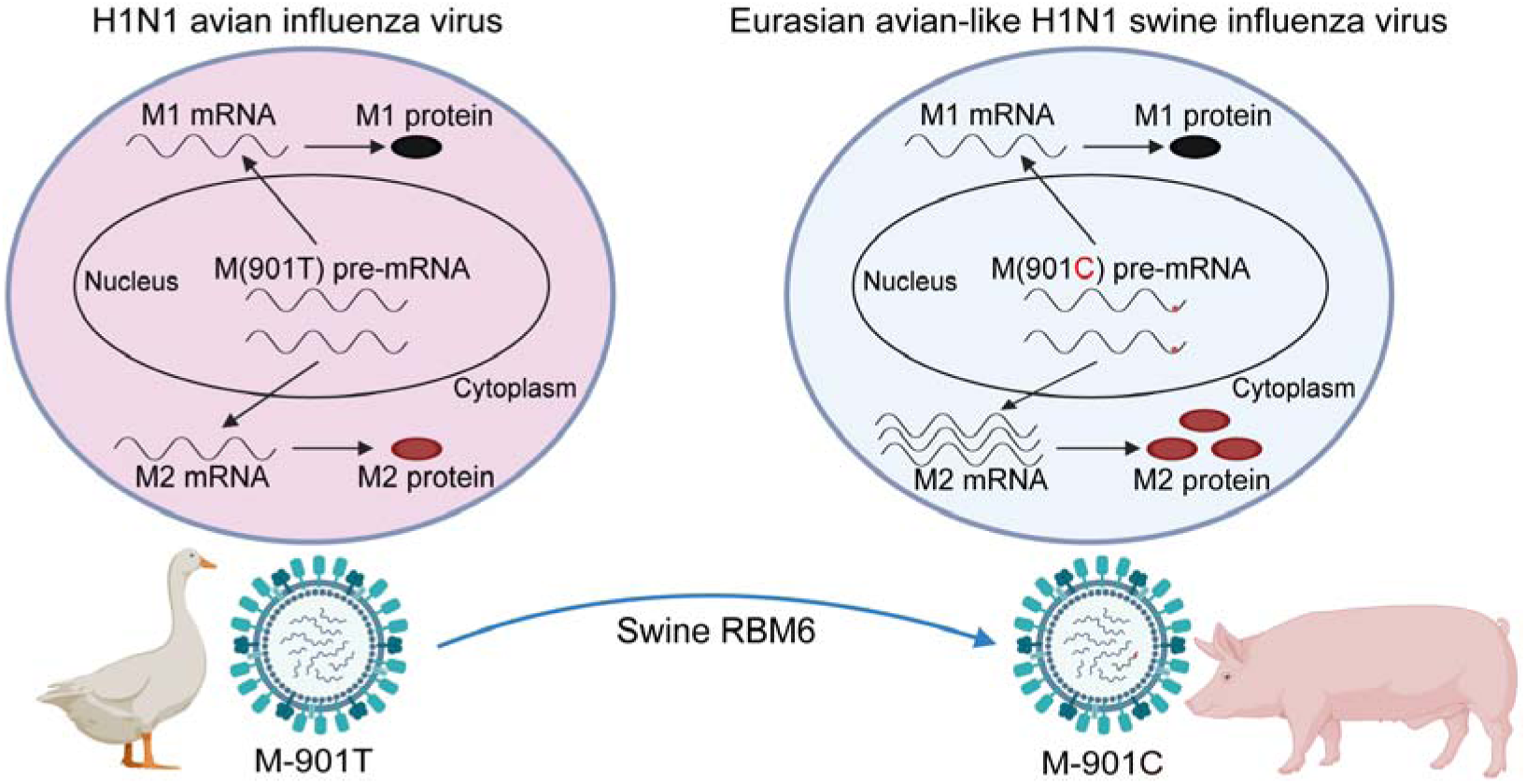
Working model of RBM6-mediated alternative splicing in facilitating the adaptation of EA H1N1 SIV. Avian influenza virus (AIV) carrying the M901T variant replicates efficiently in avian hosts, where M901 remains stable. Upon cross-species transmission to swine, swine RBM6 facilitates a nucleotide substitution from M901T to M901C. This transition enhances RBM6 binding to the viral M segment, promotes RNA splicing, increases M2 protein expression, and ultimately boosts viral replication in swine. The RBM6–M901C axis thus enables efficient adaptation of AIV to swine and supports interspecies transmission.

Our study identifies RBM6 as a critical host factor influencing influenza virus replication, yet its mode of action appears to be indirect. RBM6 belongs to the family of RNA-binding proteins but lacks intrinsic catalytic activity for RNA splicing^29^. This observation suggests that RBM6 exerts its regulatory effects by interacting with canonical splicing regulators rather than performing splicing itself. Such a mechanism is consistent with the broader paradigm in which RNA-binding proteins act as scaffolds or modulators, recruiting or stabilizing spliceosomal components to specific transcripts^30,31^. In the context of influenza virus, this implies that RBM6 may facilitate the proper processing of viral mRNAs by bridging viral RNA segments with host splicing machinery, thereby ensuring efficient production of proteins essential for replication^32,33^. Additionally, RBM6 is required for infection of EA H1N1 SIV and AIV. Interestingly, the splicing of AIV (Bav/H1N1) was not affected by RBM6-KO, suggesting that RBM6-regulated IAV infection involves alternative mechanisms. Future work should aim to delineate the precise protein-protein interactions of RBM6, as mapping its binding partners could reveal novel nodes of host-virus interplay that are amenable to therapeutic intervention.

Building on the mechanistic insights, the functionality of RBM6 appears to vary across species. Complementation assays revealed that swine RBM6 restored the replication of EA H1N1 SIV, whereas avian RBM6 failed to elicit a comparable effect. Notably, the M901C substitution was prevalent among human□infecting H1N1 IAVs, and human RBM6 similarly promoted and rescued viral replication (Supplementary Fig. 4), supporting the notion that RBM6 contributes to the host-specific adaptation of influenza viruses during replication. This divergence underscores the evolutionary adaptation of AIVs to mammalian hosts and highlights RBM6 as a potential determinant of host range. Given the low sequence homology between mammalian and avian RBM6, further comparative analysis of mammalian and avian RBM6 orthologs may identify key residues or domains that underpin functional compatibility with mammalian□adapted IAVs ^16,34^. Identifying such critical sites would not only deepen our understanding of host-specific viral replication but also provide precise molecular targets for antiviral breeding strategies in livestock ^35,36^. For example, selective modification of RBM6 alleles in pigs could reduce susceptibility to influenza virus replication, thereby mitigating zoonotic risk. This line of inquiry exemplifies how dissecting host factor differences across species can yield translational insights with direct relevance to animal health and pandemic preparedness.

Beyond host factors, our study also highlights the importance of subtle viral genetic changes in shaping replication dynamics. The mutation at position M901, involving a cytosine-to-thymine transition, represents a pyrimidine-to-pyrimidine substitution that is relatively facile from a mutational standpoint ^37,38^. Although this change does not alter the encoded amino acid, it exerts regulatory effects at the level of RNA expression. Specifically, the mutation resides within the M segment and influences the expression of M2, a protein whose role varies across influenza virus lineages. In mammalian influenza viruses, M2 appears to be more heavily utilized, whereas avian influenza viruses rely less on M2 for replication ^39,40^. This differential requirement aligns with our observation that avian-adapted viruses predominantly retain M901T, while swine-adapted viruses favor the M901C variant. Such findings suggest that synonymous mutations, often overlooked in viral evolution, can exert profound regulatory effects by modulating transcript abundance or splicing efficiency ^41,42^. The functional consequences of these mutations warrant careful attention, as they may represent hidden layers of adaptation that contribute to host specificity and pathogenicity.

In conclusion, our findings highlight RBM6 as a critical host factor that regulates influenza virus replication through modulation of M segment splicing, with the M901C site serving as a key determinant of swine adaptation. The RBM6-M901 axis represents a previously unrecognized mechanism of viral evolution and interspecies transmission. Surveillance of M901 variation across influenza subtypes, coupled with functional studies of RBM6 interactions, may provide early warning of strains with enhanced pandemic potential and inform strategies for therapeutic intervention.

## Materials and Methods

### Ethics statement

Approval for the animal experiments carried out in this study was obtained from the Committee on the Ethics of Animal Experiments at Huazhong Agricultural University (No. HZAUMO-2025-0106).

### Cells

Human embryonic kidney 293T cells (HEK293T, Cat# CRL-3216), Madin-Darby canine kidney (MDCK, Cat# CCL-34) cells, Porcine Kidney-15 (PK-15, Cat# CCL-33) cells, UMNSAH/DF-1 cells (DF-1, Cat# CRL-3586), and adenocarcinomic human alveolar basal epithelial cells (A549, Cat# CCL-185) were acquired from the American Type Culture Collection (ATCC, Manassas, VA, USA). Stably expressing Cas9 of PK-15 (PK-15-Cas9) and A549 (A549-Cas9) cell lines were established through selection via puromycin or blasticidin. All cell lines were cultured at 37°C in a humidified atmosphere containing 5% CO_2_, utilizing RPMI 1640 (SH30809.01, HyClone, USA) or Dulbecco’s Modified Eagle Medium (DMEM) (SH30243.01, HyClone, USA), supplemented with 10% fetal bovine serum (FBS) (FSP500, ExCell, China).

### Viruses and reverse genetics

The influenza A viruses (IAVs) employed in this study included A/duck/Bavaria/2/1997 (Bav/H1N1), A/Swine/HuBei/221/2016 (HuB/H1N1), and A/Hunan/42443/2015 (HuN/H1N1), and A/Puerto Rico/8/1934 (PR8/H1N1). Recombinant viruses were constructed using the genetic background of A/Hunan/42443/2015 (HuN/H1N1) through an eight-plasmid reverse genetic system ^15^. Amplification of all other viruses was conducted in 10-day-old embryonated chicken eggs, with titration performed by determining the TCID_50_ values in MDCK cells.

### Plasmids

Lentiviruses were produced using the lenti-sgRNA-EGFP and lenti-guide-puro vectors, along with the pMD2.G and psPAX2 plasmids ^43^. For the construction of the lentiviral sgRNA vector, paired sgRNA oligonucleotides (50 μM per oligo) were annealed and cloned into lenti-sgRNA-EGFP or lenti-guide-puro vector, which was linearized with *Bbs*I or *Bsm*BI (R3539 and R0739, NEB, USA). The pCAGGS-HA-RBM6 (HA-RBM6) of different species constructed by cloning the full-length cDNA into the pCAGGS-HA vector, respectively. Eight segments of A/Hunan/42443/2015 (HuN/H1N1) were inserted into the pHW2000 vector, and mutant M genes targeting base 901 were generated by PCR-based site-directed mutagenesis, confirmed by sequencing ^44^.

### Antibodies and reagents

The antibodies and reagents used in the study were as follows: Rabbit anti-RBM6 (14360-1-AP, Proteintech, China); mouse anti-HA tag (66006-2-Ig, Proteintech, China); mouse anti-glyceraldehyde-3-phosphate dehydrogenase (GAPDH) (60004-1-Ig, Proteintech, China); rabbit anti-IAV NP, M1, M2, NS1, and NS2 (GTX125989, GTX125928, GTX125951, GTX125990, and GTX125953, GeneTex, USA); horseradish peroxidase-conjugated goat anti-mouse and anti-rabbit (AS003 and AS014, ABclonal, China); Cy3-conjugated Goat anti-Rabbit IgG (H+L) (AS007, ABclonal, China), and 4′,6′-diamidino-2-phenylindole (1:5,000) (C1002, Beyotime, China).

### Generation of RBM6 knockout cell line using CRISPR/Cas9

Individual sgRNA constructs targeting RBM6 were generated and incorporated into the lenti-sgRNA-EGFP or lenti-guide-puro vector. Lentiviruses were produced following established protocols ^45^. These lentiviruses were then transduced into PK-15-Cas9, DF-1, or A549-Cas9 cells, respectively. Transduced cells were selected using fluorescence-activated cell sorting (FACS) or 1.5 μg/mL puromycin. Monoclonal cells were obtained through the limiting dilution method and subsequently expanded. Confirmation of RBM6 knockout cells was achieved through Sanger sequencing and western blot analysis.

### Cell viability assay

To examine the impact of RBM6-KO on cellular proliferation, the viability of RBM6-KO cells and WT cells was measured through CCK-8 activity, following the manufacturer’s instructions. Briefly, cells were seeded onto 96-well plates, and their viabilities were measured at 12-, 24-, and 36-hours post-seeding. CCK-8 reagent (CK04-500T, Dojindo Molecular Technologies, Japan) was applied to each well. and the subsequent measurement of absorbance at 450 nm was conducted using a microplate reader after a 1-hour incubation at 37°C in dark.

### Transfection

Transfections were performed using Lipofectamine 2000 (11668019, Invitrogen, USA) according to the manufacturer’s instructions. Briefly, plasmids and Lipofectamine were diluted to equal volumes with Opti-MEM and incubated for 5 min at room temperature. The diluted Lipofectamine and plasmids were mixed and incubated for 20 min at room temperature. The mixture was added to cells and incubated for 6 h, and the medium was then replaced with fresh medium supplemented with 10% FBS.

### Virus infection and titration

To evaluate the impact of RBM6 knockout on viral replication, wild type and RBM6-KO cells were independently seeded in triplicate within 12-well plates. For IAV infection, cells underwent two times washes with DMEM, followed by incubation with diluted virus at the MOI of 0.01 for 1 hour. Subsequently, cells were again washed twice with DMEM and replenished with fresh infection medium (DMEM supplemented with 0.2 μg/mL TPCK-treated trypsin (T1426, Sigma-Aldrich, USA)). Supernatants were collected at designated time points post-infection, and viral supernatants were serially diluted with DMEM. Eight replicates of each dilution were added to the wells, and the 50% tissue culture infectious dose (TCID_50_) was calculated using the Reed-Muench method 72 hours after infection ^46^.

### Indirect immunofluorescence assay and confocal microscopy

WT and RBM6-KO PK-15 cells were incubated with HuB/H1N1 at an MOI of 0.1 for 1 h. The cells were fixed with 4% paraformaldehyde (PFA) for 10 min at indicated timepoints, followed by incubating with 1% (wt/vol) bovine serum albumin (BSA) for 1 h at room temperature. Samples were then incubated with the anti-NP antibody (GTX125989, GeneTex, USA) for 2 h at room temperature, followed by incubation with the Cy3-conjugated secondary antibody for 1 h. The nuclei were stained with DAPI for 10 min at room temperature. Images were acquired using a confocal microscope (LSM880, Zeiss, Germany).

### RNA isolation and qRT-PCR

For quantitative reverse transcription-PCR (qRT-PCR), cells were lysed with TRIzol reagent (15596018, Invitrogen, USA), and total RNA was extracted according to the manufacturer’s instructions. 1 μg of RNA was used to generate cDNA using reverse transcriptase (RK20403, ABclonal, China). Real-time PCR (Vii7A; ABI, USA) was performed using SYBR GREEN (RK21203, ABclonal, China). The PCR conditions were 2 min at 50°C, 10 min at 95°C, and then 40 cycles of 5 s at 95°C and 30 s at 60°C. GAPDH mRNA was used as a control for the normalization of cellular and viral mRNA.

### CLIP-Seq

Stably expressing halo-tagged RBM6 cells were infected with HuB/H1N1 at an MOI of 5 for 9 h, followed by irradiation with UV light at 365 nm to induce covalent crosslinking between RBM6 and its bound RNAs. Cell lysates were prepared and subjected to immunoprecipitation using halo-specific antibodies conjugated to magnetic beads (G7281, Promega, USA), thereby enriching RNA–protein complexes. The complexes were digested with proteinase K to remove protein components, and the recovered RNA fragments were processed by end repair and adaptor ligation. Reverse transcription and PCR amplification were performed to generate sequencing libraries. High-throughput sequencing was conducted on an Illumina platform, and bioinformatic analysis was used to align reads to the reference genome and identify binding sites within viral segments ^22,25^.

### RNA pull-down

Biotin-labeled RNAs were either synthesized by Tsingke or generated via *in vitro* transcription (6140, Takara Beijing, China), and subsequently modified using the Pierce RNA 3′ End Desthiobiotinylation Kit (20163, Thermo Fisher, USA). Cell lysates were prepared in RIP buffer (20 mM Tris-HCl, pH 7.5; 150 mM KCl; 0.5% NP-40; 2 mM EDTA; protease inhibitor cocktail; RNase inhibitor). Biotinylated RNAs were pre-bound to streptavidin magnetic beads and incubated with lysates at 4°C overnight according to the manufacturer’s instructions (20164, Thermo Fisher, USA). After extensive washing, bound proteins were eluted and analyzed by SDS-PAGE followed by immunoblotting with specific antibodies.

### RNA immunoprecipitation (RIP) assay

Wild-type PK-15 cells were infected with either wild-type or the M C901T mutant HuN/H1N1 for 12 h. Following infection, RIP was performed using anti-RBM6 antibody (14360-1-AP, Proteintech, China) or control IgG (AC005, ABclonal, China). The immunoprecipitated RNA was extracted using a standard RNA isolation protocol, and the abundance of viral M segment RNA was quantified by qRT-PCR ^47^.

### *In vivo* experiments

For an in-depth exploration of the role of RBM6 and viral M 901C in EA H1N1 SIV infection *in vivo*, RBM6 knockout mice were generated by CRISPR-Cas9. 6-week-old WT and RBM6^+/-^ C57BL/6N mice were nasally challenged with WT or C901T mutant HuN (H1N1), respectively. Daily monitoring of body weight loss and survival occurred over 2 weeks post-infection (n = 10). Mice exceeding a 30% loss in initial body weight were humanely euthanized ^21,48^. On 3 and 5 days post-challenge, a subset of mice from each group (n = 3) underwent anesthesia, and sacrifice and their lungs were either homogenized and/or fixed in 4% formaldehyde. The homogenized lung samples were utilized for assessing gene expression as well as virus titers. The fixed mouse lung samples were used for hematoxylin & eosin (H&E) and immunofluorescence staining for histopathological analysis.

### Statistical analysis

All measurements were taken in triplicate, and the presented data are outcomes from at least two separate experiments. The results are shown as the mean ± standard deviation of the triplicate determinations. Statistical significance was ascertained by computing *P* values using the paired two-tailed Student’s t-test (ns, *P* >0.05; *, *P* < 0.05; **, *P* < 0.01; ***, *P* < 0.001; ****, *P* < 0.0001).

## Data availability

All data are available in the Article and its Supplementary Information.

## Acknowledgments

We would like to thank the National Key Laboratory of Agricultural Microbiology Core Facility for assistance in confocal microscopy and flow cytometry. This work was supported by the National Natural Science Foundation of China (32430104 and 32025036 to H.Z.), the National Key Research and Development Program (2024YFE0106100 to J.Z. and W.S.), the Fundamental Research Fund for the Central Universities (2662025DKPY009 to H.Z.), Hubei Hongshan Laboratory (2022hszd005 to H.Z.), and the earmarked fund for CARS-41 to H.Z.

## Author contributions statement

H.Z. conceived the project; J.Z., S.T., H.S., C.X., M.J., J.G., and S.T. conducted the experiments; J.Z., S.T., H.S., T.C., T.P.P., W.S., W.S.B., and H.Z. analyzed the data J.Z. and H.Z. wrote and revised the paper. All authors reviewed and approved the final manuscript.

---

**Supplementary Fig. 1.**
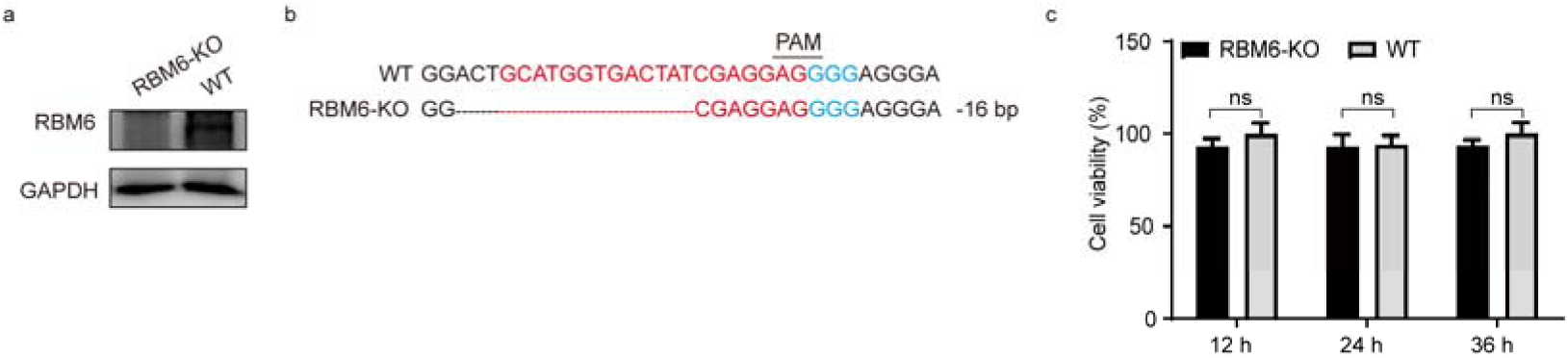
Determination of the knockdown efficiency of RBM6 and its effect on cell viability. (a) Western blot analysis showing the expression of the RBM6 protein in RBM6-KO and WT PK-15 cells, with GAPDH gene as an endogenous control. (b) Sanger sequencing confirmation for the generated RBM6-KO cell lines using CRISPR/Cas9 technology. The sgRNA sequence was highlighted in red and NGG sequence was highlighted in blue. PAM: Protospacer Adjacent Motif. (c) Cell viability in RBM6-KO and WT PK-15 cells was determined using CCK-8 reagents over 36 hours. Error bar in panel c indicates the standard deviation. The data shown in panel c is means□±□SD (n□=□8 biologically independent experiments). Statistical analysis was performed using unpaired, two-tailed Student’s *t*-test. (ns, *P* > 0.05).

**Supplementary Fig. 2.**
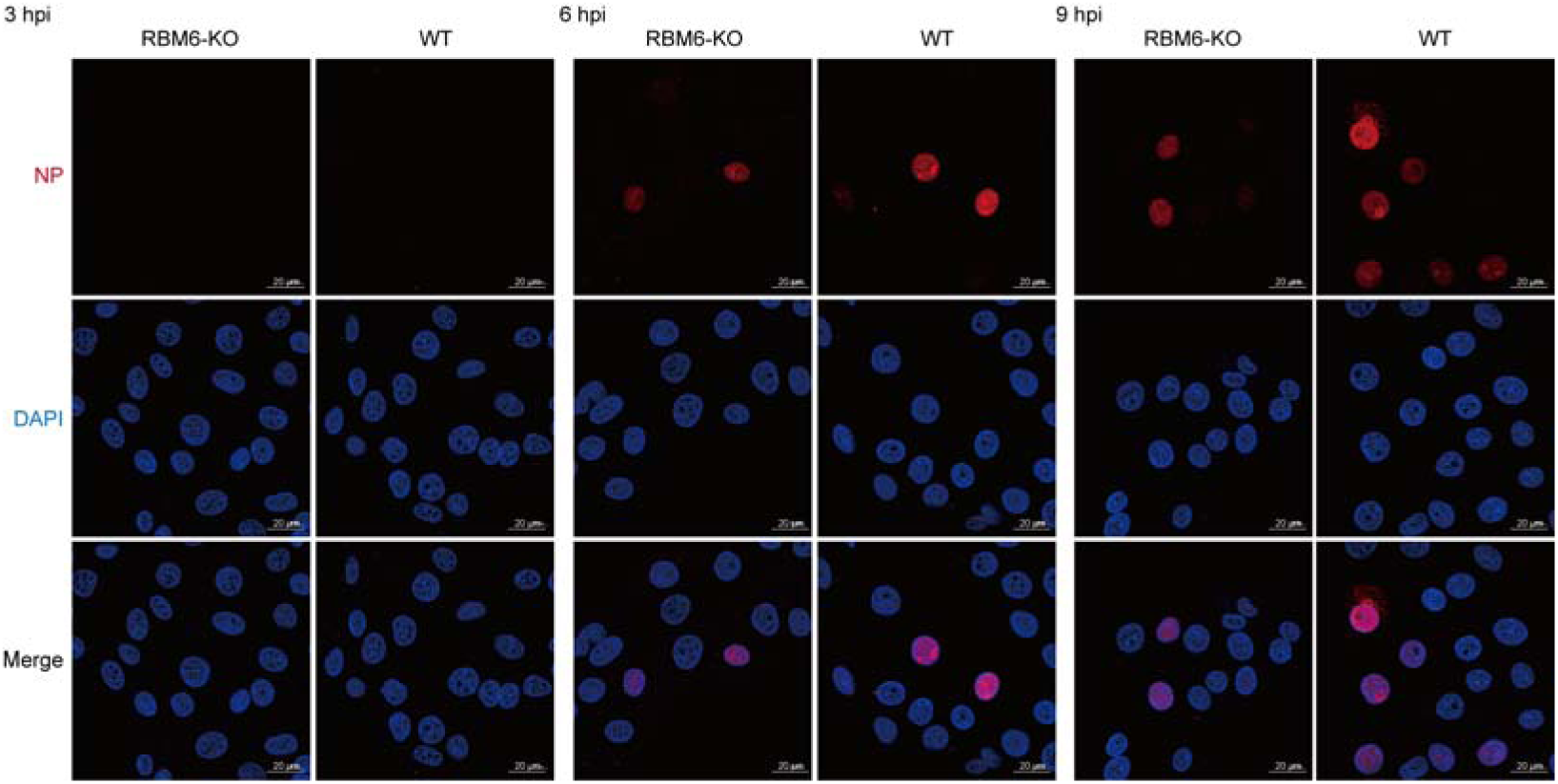
RBM6 is involved in the EA H1N1 SIV replication after nuclear import. RBM6-KO and WT PK-15 cells were infected with HuB/H1N1 (MOI = 0.1). Cells were fixed at 3, 6, and 9 hpi, and cells were stained for viral NP (red) using the anti-NP antibody. DAPI was used to stain for the nucleus (blue). Scale bars = 20 μm.

**Supplementary Fig. 3.**
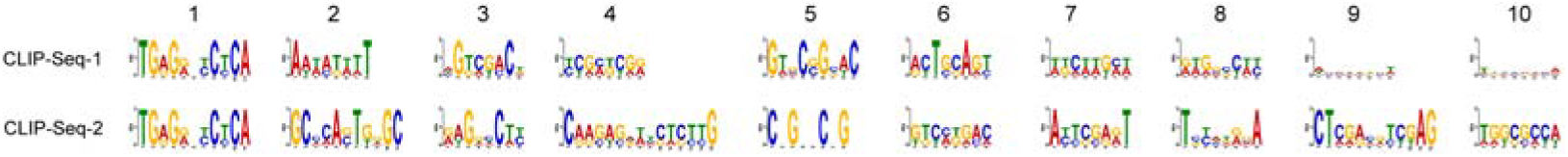
Top binding motifs of RBM6 identified on viral M and NS segments. RBM6-binding motifs were screened and ranked according to their enrichment across the viral M and NS gene segments. The analysis highlights the most conserved and high-affinity motifs.

**Supplementary Fig. 4.**
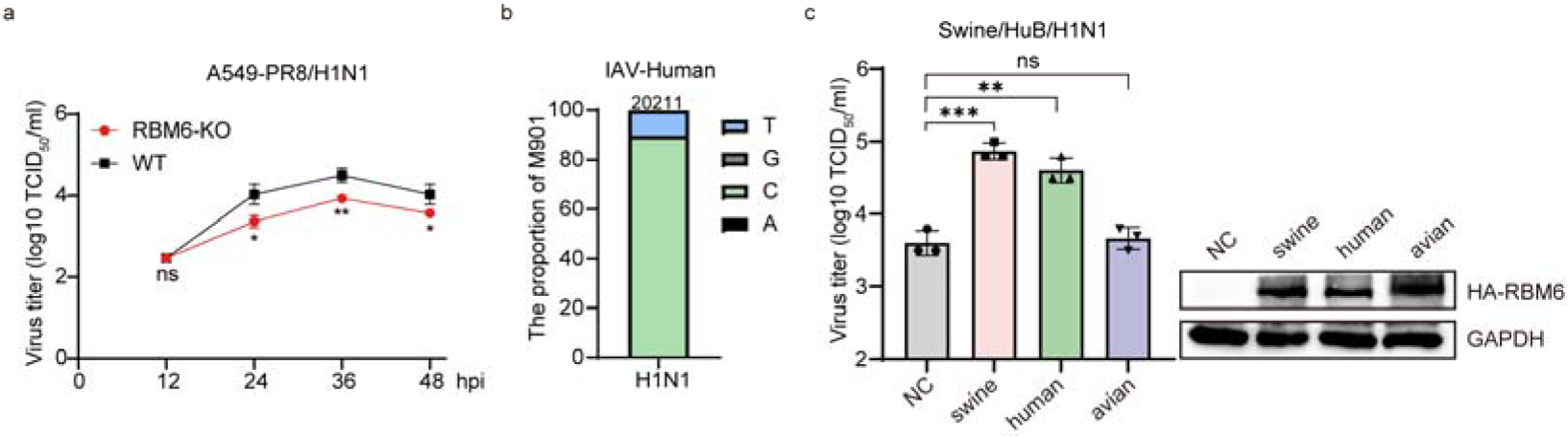
Human RBM6 promoted the replication of IAV. (a) Effect of RBM6 knockout in A549 on the replication of IAV. RBM6-KO and WT A549 cells were infected PR8/H1N1 at a MOI of 0.01, and the viral titers at indicated timepoints were determined by TCID_50_. (b) Conservation analysis of the M901 site among human-infecting H1N1 IAVs downloaded from the National Center for Biotechnology Information (NCBI). (c) Restoration of mammalian RBM6 promotes replication of EA H1N1 SIV. Exogenous RBM6 (HA RBM6) from different species was transfected into RBM6□KO PK□15 cells. Transfected cells were infected with HuB/H1N1 at an MOI of 0.1, and viral titers at 24 hpi were determined by TCID_50_. Corresponding protein expression was detected by western blot. Error bar indicates the standard deviation. The data shown in panels a-b are means□±□SD (n□=□3 biologically independent experiments). Statistical analysis was performed using unpaired, two-tailed Student’s *t*-test. (ns, *P* > 0.05; *, *P* < 0.05; **, *P* < 0.01; ***, *P* < 0.001).

